# Detecting Full-Length EccDNA with FLED and long-reads sequencing

**DOI:** 10.1101/2023.06.21.545840

**Authors:** Fuyu Li, Wenlong Ming, Wenxiang Lu, Ying Wang, Xiaohan Li, Xianjun Dong, Yunfei Bai

## Abstract

Reconstructing the full-length sequence of extrachromosomal circular DNA (eccDNA) from short sequencing reads has proved challenging given the similarity of eccDNAs and their corresponding linear DNAs. Previous sequencing methods were unable to achieve high-throughput detection of full-length eccDNAs. Here we describe a new strategy that combined rolling circle amplification (RCA) and nanopore long-reads sequencing technology to generate full-length eccDNAs. We further developed a novel algorithm, called Full-Length eccDNA Detection (FLED), to reconstruct the sequence of eccDNAs. We used FLED to analyze seven human epithelial and cancer cell line samples and identified over 5,000 full-length eccDNAs per sample. The structures of identified eccDNAs were validated by both PCR and Sanger sequencing. Compared to other published nanopore-based eccDNA detectors, FLED exhibited higher sensitivity. In cancer cell lines, the genes overlapped with eccDNA regions were enriched in cancer-related pathways and *cis*-regulatory elements can be predicted in the upstream or downstream of intact genes on eccDNA molecules, and the expressions of these cancer-related genes were dysregulated in tumor cell lines, indicating the regulatory potency of eccDNAs in biological processes. Our method takes advantage of nanopore long reads and enables unbiased reconstruction of full-length eccDNA sequences. FLED is implemented using Python3 which is freely available on GitHub (https://github.com/FuyuLi/FLED).

## 1 Introduction

Extrachromosomal circular DNA is a class of chromosomal original DNA with a covalently closed circular structure which was found common and highly conversed in eukaryotes ranging from yeast to human [1-3], with lengths ranging from tens to millions of base pairs [4, 5]. eccDNAs arising from unique non-repetitive genomic regions have been discovered in normal and malignant cells [6, 7], which carry intact genes and specific regulatory elements [6-8]. The existence of eccDNA will lead to genomic instability while its regulatory capacity could contribute to aging [9] and intercellular genetic heterogeneity in various tumors [10]. By amplifying oncogenes [11-13] or therapeutic resistance genes [14, 15] located on it, eccDNAs can drive genome remodeling [16] and distal regulation [7, 17]. To effectively purify and amplify eccDNA for de novo detection, a sensitive method called Circle-seq [18, 19] has been applied followed by high-throughput sequencing. Briefly, this approach encompasses column purification, removal of remaining linear chromosomal DNA by exonuclease, and rolling-circle amplification (RCA). Several computational methods were developed to identify eccDNA from discordantly mapped paired-end reads as well as soft clipped reads around eccDNA breakpoint [3, 20]. However, limited by the relatively short length of Illumina sequencing reads, these methods were restricted to detect reads that span the eccDNA breakpoints and therefore cannot determine the internal structure of the eccDNA, especially for eccDNAs including parts from different chromosomes [11, 21]. The function of eccDNA is fundamentally determined by its constituent sequence elements, hence obtaining the full sequence of eccDNA is essential for understanding eccDNA structure, transcription, and regulatory function. Although the sequence of eccDNA can be inferred by reconstructing the internal structure of the DNA amplifications [11, 21], the method of assembling short-read data inevitably introduces errors, especially for homologous or similar eccDNA molecules. Long-reads-based sequencing methods like Oxford Nanopore [22] have the advantage of real-time ultra-long read sequencing, allowing to cover the sequence of the whole molecule in one read. As we noticed, there are few existing bioinformatics tools to identify eccDNA from RCA nanopore data: CReSIL [23], cyrcular-calling-work-flow [24], Nanocircle [25], eccDNA_RCA_nanopore [26, 27], and ecc_finder [28]. Similar to short-reads-based methods, Nanocircle determined the joined breakpoint coordinates of genomic regions during eccDNA fragment formation by exploiting split reads while CReSIL reconstructed the sequence of eccDNA fragments by de novo assembly of sequence reads. The cyrcular-calling-workflow constructed a directed graph based on read depth information and split reads, and then called eccDNAs by calculating the posterior probability for each plausible circular path[24]. The eccDNA_RCA_nanopore method detected full-length eccDNAs by bootstrapping successive aligned subreads in each RCA long read [26], while ecc_finder identified candidate reads with a tandem repeat pattern and divided each read into repeat subread first and then determined the fragment composition of full-length eccDNA by the genomic mapping locations of the repeat subread [28]. Therefore, ecc_finder and eccDNA_RCA_nanopore could obtain the full-length sequences of eccDNAs directly from sequencing reads without assembly or splicing, but both required supporting reads overriding the joint break-point of original eccDNA at least twice to ensure the repeat patterns in RCA reads, which is severely limited by the size of the eccDNAs and the amplification enzyme activity, resulting in a poor utility of the sequencing data.

To address this challenge, we have developed a new bioinformatics method, called **F**ull **L**ength **E**ccDNA **D**etection (FLED), taking the advantage of the nanopore long-read sequencing technology and rolling-circle amplification method, to profile full-length eccDNAs effectively. The FLED process includes three steps. Candidate eccDNA detection was designed to construct two sets of weighted directed graphs based on the split alignment of each RCA long read: the first one aimed to find the optimal successive alignment of each read according to the location of every subread on it, and the second one was constructed based on the genomic coordinates of every subread in the optimal alignment and the candidate eccDNAs were identified by closed-cycle detection. For a given eccDNA candidate, all supporting reads were cut at the breakpoints, and one high-quality consensus sequence was produced as the full-length sequence for each candidate. Finally, the number of supporting reads and the variation of sequencing depth around the joined breakpoint were calculated and used to filter eccDNAs with high confidence. The adapted Circle-seq and FLED workflow is summarized in Figure 1.

**Figure 1.**
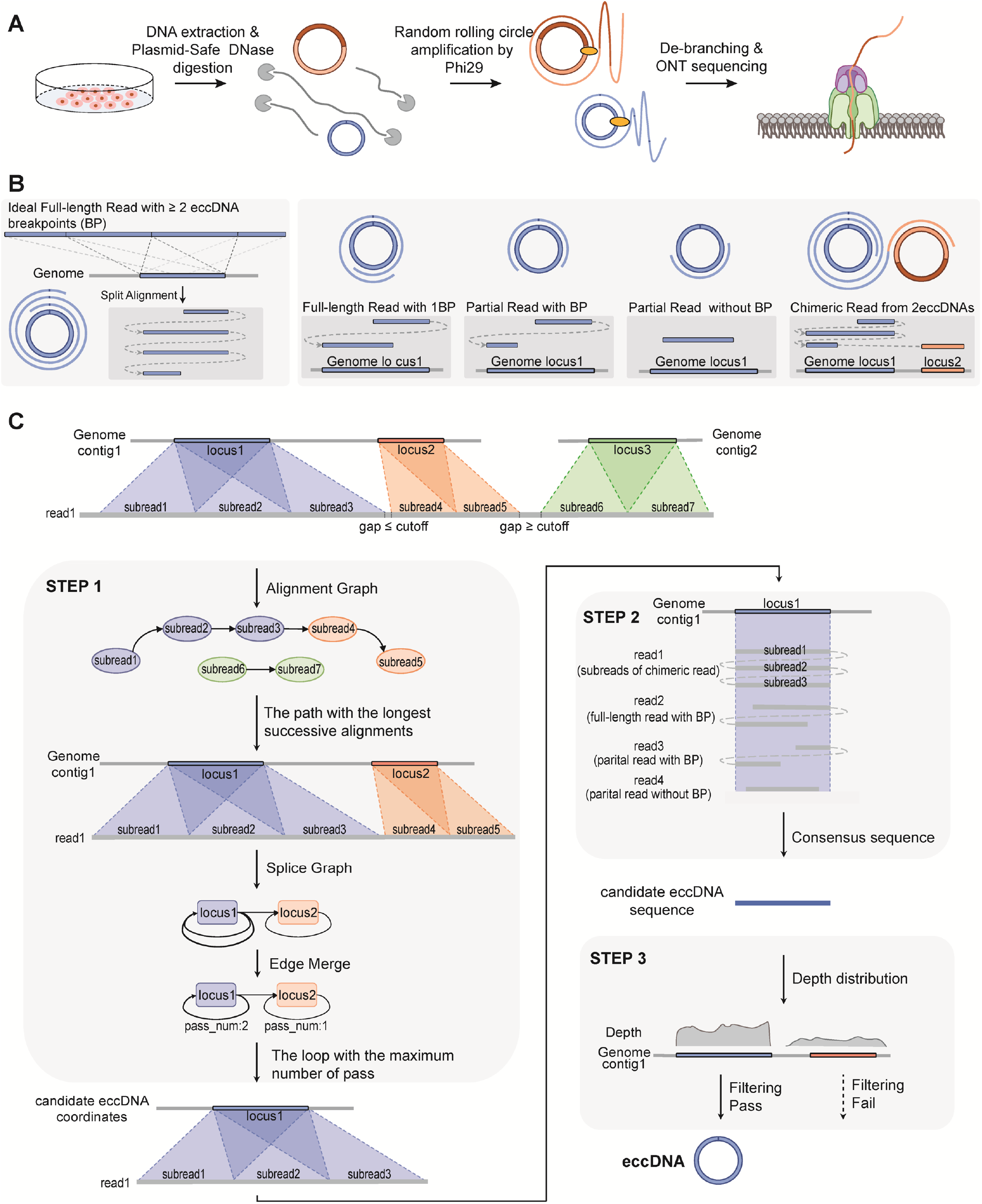
Workflow of adapted Circle-seq and FLED. **A**, Construction of nanopore sequencing libraries. The genomic DNA was extracted and subjected to Plasmid-Safe DNase digestion to degrade linear dsDNA. The remaining DNA was performed by random rolling circle amplification by Phi29. Product DNA was de-branched and sequenced using the MinION platform. **B**, Diagram showing the different types of RCA products. For eccDNA from a single genomic locus, the ideal RCA long reads should cross the joined breakpoints at least twice, so the continuous subreads were aligned to a single locus (left panel). Reads covered the joined breakpoint once can be classified as full-length reads or partial reads based on the overlapping of subreads. Partial reads covering breakpoints miss the internal structure and partial reads without breakpoints miss the genomic start and end position of eccDNA. **C**, Workflow of eccDNA identification for aligned high-quality nanopore reads. STEP 1: Candidate eccDNA detection. STEP 2: Full-length sequences correction. STEP 3: False-positive reduction.

## 2 Materials and Methods

### 2.1 EccDNA extraction and sequencing

In this study, we studied eccDNAs from seven human cell lines: gastric cancer cells BGC823, gastric cancer cells SGC7901, gastric epithelial cells GES1, hepatocellular carcinoma cells HepG2, liver cells HL7702, breast carcinoma cells MDA-MB-453 (MB453), and breast epithelial cell line MCF12A. Each sample was subjected to Plasmid-Safe ATP-dependent DNase (Epicentre) digestion and performed rolling circle amplification by Phi29 DNA polymerase with Exo-resistant random primer followed by Nanopore sequencing according to the manufacturer’s protocol (**Figure 1A**, Detailed in **Supplementary Methods**). High-quality base-called reads were aligned to the human reference genome (GRCh38) using long-read splice-aware mapper minimap2 [29, 30], and then input for eccDNA detection using our FLED method.

### 2.2 Full-length eccDNA detection method

We developed a novel graph-based algorithm for **f**ull-**l**ength **e**ccDNAs **d**etection, called FLED. FLED takes long-read sequencing data or splice-aware alignment as input and identifies full-length eccDNAs in three main steps (**Figure 1B-C**).

#### 2.2.1 Candidate eccDNA detection

As minimap2 does split-read alignment, nanopore ultra-long reads with high alignment quality were split into multiple aligned subreads. To remove misalignments of subreads, a weighted directed alignment graph based on the order of aligned subread was constructed for each RCA long read with nodes representing aligned subreads and the edge indicating the adjacency of two subreads on raw read. The length of the path in the alignment graph was calculated as the total length of all subreads on this path and the set of subreads on the longest path represented the optimal successive alignment of this RCA long read. To reduce the impact of possible DNA contamination during the RCA process, FLED set up a hard filter on the alignments, where the total length of consecutive aligned subreads exceeded 70% of the read length. For filtered reads with only two subreads, FLED identified eccDNAs and classified them according to the overlap of genomic mapping positions of these two subreads (**Figure 1B**). For filtered read with more subreads, FLED builds a splice graph based on the genomic coordinates of each subread to identify eccDNAs. Each splice graph is a directed graph in which nodes represent genomic fragments while the weighted directed edges represent the number of connections between two such nodes based on their adjacent parts. Theoretically, the splice graph should be a closed circuit owing to the nature of eccDNAs and the closed traversal path represents the composition of a candidate full-length eccDNA. FLED finds a closed traversal path with the maximum total weight as the composition of the eccDNA by depth-first search (DFS). Similarly, FLED discards chimeric amplification product of the original eccDNA molecule where the total length of all subreads in the loop is less than 50% of raw read, due to possible template-dependent polymerase jumping occurring during RCA.

Furthermore, resulting from the large size of few eccDNAs and the low efficiency of RCA, over 90% of nanopore reads contained only partial sequences of eccDNAs. To improve data utilization, FLED identifies eccDNAs from reads that crossed joined breakpoints (**Figure 1B**).

#### 2.2.2 Full-length sequences correction

The sequence of eccDNA can be completely preserved in the RCA long read, but at the same time, the inaccuracy of nanopore sequencing technology also leads to the failure of using the raw sequence of the subread directly as the sequence of eccDNA. To avoid the uncertainty of assembly, the correction was performed on the full-length eccDNAs only, which was supported by at least one full-length read. FLED truncates uncorrected full-length reads into subreads according to the closed-loop detected in the splice graph and groups all subreads based on the putative eccDNAs they support. To improve the accuracy, FLED incorporates high-quality non-full-length reads mapped to the same genomic region and generates consensus sequences for each group as accurate full-length sequences of Full eccDNAs by partial order alignment algorithm SPOA [31].

#### 2.2.3 False-positive reduction

FLED implements hard filters at the alignment and eccDNA level to reduce eccDNA identification errors. At the alignment level, FLED examines the mapping qualities and alignment length as described while at the eccDNA level, FLED applies three filters to the detected eccDNAs. Real eccDNAs are resistant to exonuclease digestion and are supposed to show high sequencing depth inside the junction. Therefore, the candidate eccDNA supported by at least 2 distinct nanopore reads was further validated by the sequencing depth adjacent to its joined breakpoints. For each candidate eccDNA, FLED calculates the mapping depth for 50 bp upstream to 50bp downstream around its breakpoints and outputs the candidate when two criteria were met: the ratio between the inside depth and overall depth should be greater than 0.5 while the variation of sequencing depth between inside and outside should be significant based on a Wilcoxon test (P < 0.05). All the parameters for reducing false positives are adjustable to adapt to various data and requirements.

### 2.3 EccDNA validation by PCR and Sanger sequencing

To verify the junction sequences of eccDNAs, a pair of PCR primers specific for breakpoint junctions were designed using Primer-Premier 5 (Premier Biosoft Interpairs, Palo Alto, CA). The outward PCR experiments were performed using NEBNext High-Fidelity 2X PCR Master Mix (NEB). For a 50 µl reaction, 10 ng of the MDA products were used, followed by an initial denaturing step of 98 °C for 30 s; 35 cycles of 98 °C for 10 s, 60 °C for 20 s, 72 °C for 20 s; and a final extension of 72 °C for 2 min. The PCR products were analyzed by agarose gel electrophoresis and performed Sanger sequencing. Primer sequences and expected PCR product sizes of the split junction sequences are detailed in **Supplementary Table 1**.

## 3. Results

### 3.1 EccDNA detection and validation in simulated datasets

FLED was evaluated in simulated datasets first. Both high- and low-coverage datasets were simulated by NanoSim [32] (Detailed in **Supplementary Methods**), resulting in 4,297 and 4,153 simulated eccDNAs under the sequencing depths of 30x and 10x, respectively. We then compared the FLED-identified eccDNAs with simulated eccDNAs to evaluate the overall performance of FLED with a single metric F1 score. On high-coverage data, FLED attained an F1 score of 0.97 by detecting 4,058 out of the 4,297 simulated eccDNAs, while moderate sensitivity and precision were also found on low-coverage datasets with an F1 score of 0.77. With increased sequencing depth, the proportion of full-length eccDNA raised from 53.81% to 72.95%. The length distribution of full-length sequences obtained by FLED displayed a high correlation with the simulated length (Pearson’s r = 0.9, p-value of 2.2×10^−26^) (**Supplementary Figure 1**). These results indicated that sequencing depth plays a significant role in eccDNA detection, especially for full-length eccDNA. More importantly, FLED still performed well on low-depth data.

### 3.2 Verifying eccDNAs by PCR, Sanger sequencing, and comparison

To validate the accuracy of eccDNA detection by FLED, RCA nanopore sequencing was applied for the GES1 cell line and 5780 eccDNAs were detected based on FLED, and 11 eccDNAs of different sizes ranging from 600 to 9,600 bp were randomly selected and the outward primers were designed to amplify the junction region. 90% (10 of 11) selected eccDNA were successfully validated by both PCR and Sanger Sequencing (**Figure 2A**).

**Figure 2.**
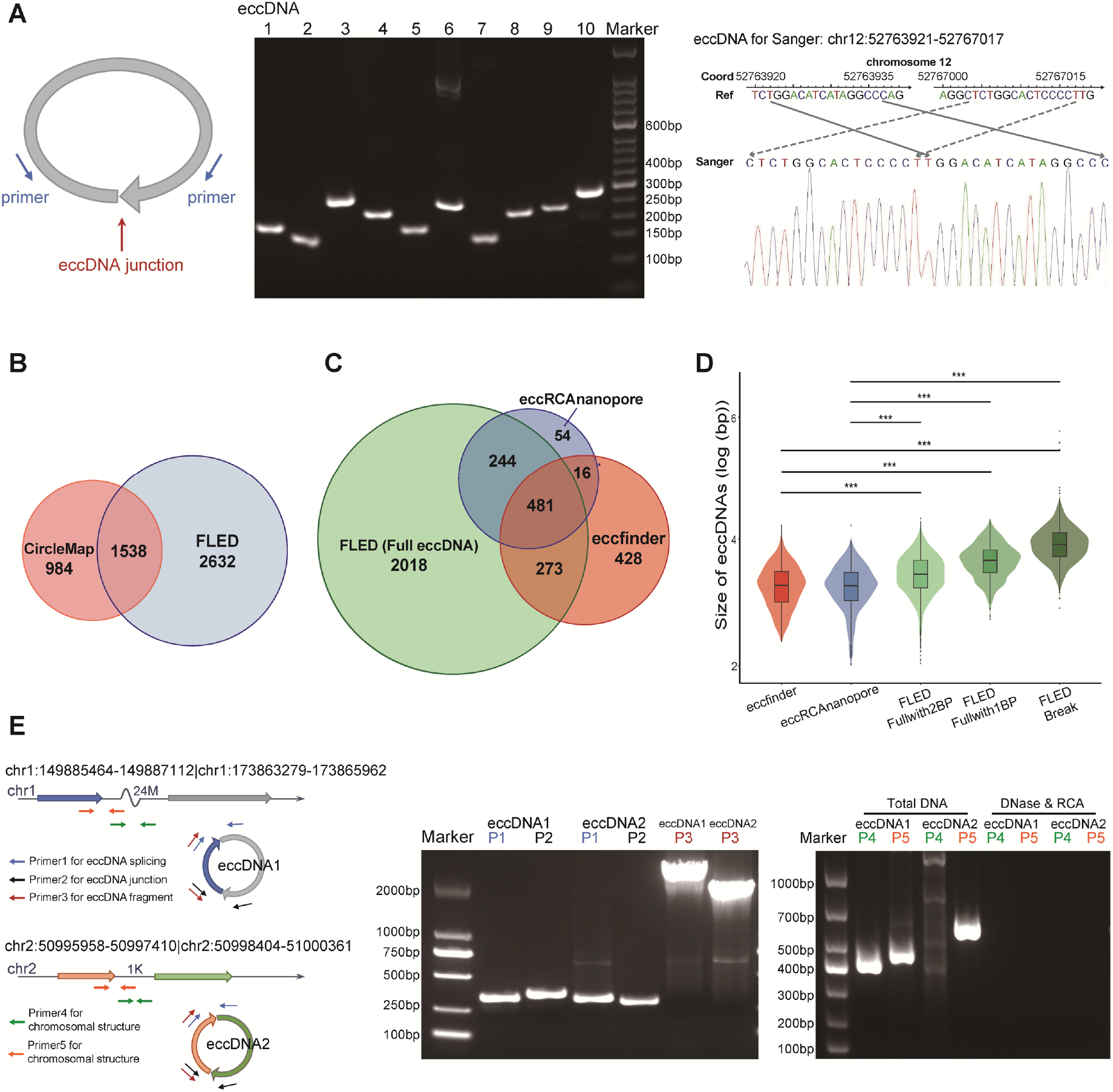
eccDNAs validation and comparison in GES1 cell line. **A**, Gel images for a validated subset (n = 10) of eccDNAs by outward PCR and the Sanger sequencing of outward PCR product of validated eccDNA example. EccDNAs are named according to the coordinates of breakpoints (red arrow). The list of validated eccDNAs and corresponding primer (blue arrow) are summarized in **Supplementary Table 1. B**, The eccDNA detected by Circle-Map (2,522 in total) for NGS data and FLED (5,780 in total) for Nanopore data, and identical eccDNA (1,538) detected by both methods. **C, D**, Comparison of three nanopore-based eccDNA detectors. **C**, The number of full-length eccDNAs detected by ecc_finder (1,198 in total), eccDNA_RCA_nanopore (795), and FLED (3,016). **D**, The size distribution of detected eccDNAs. eccfinder: eccDNAs detected by ecc_finder; eccRCAnanopore: eccDNAs detected by eccDNA_RCA_nanopore; FLED Fullwith2BP: FLED-detected Full eccDNAs supported by reads overriding the joined breakpoint at least twice; FLED Fullwith1BP: FLED-detected Full eccDNAs supported by reads overriding the joined breakpoint only once; FLED Break: Break eccDNAs detected by FLED. P-values are determined using the Wilcox test. Significance: (***) P-value<0.001; **E**, Gel images for 2 multiple-fragments eccDNAs by PCR. The list of validated eccDNAs and corresponding primer sequences are summarized in **Supplementary Table 1**.

The Illumina library was constructed for the GES1 cell line and sequenced for comparison and validation. Circle-Map was deployed for eccDNA detection on NGS data and the detected eccDNAs by Circle-Map were **filtered under the same criteria as FLED used. Finally, a total of 2**,**522** eccDNAs breakpoints were detected by Circle-map, while 5,780 by FLED including 4,170 Full eccDNAs. We found that 1,538 eccDNAs overlapped in both results, accounting for 62% and 27% of Circle-map and FLED respectively (**Figure 2B**).

We further compared the performance of FLED with two other published full-length eccDNA detectors based on nanopore data which both identify full-length eccDNA from full-length reads crossing the joined breakpoints as least twice. In the GES1 cell line, 795, 1198, and 3016 eccDNAs with at least 3 supporting reads were identified by eccDNA_RCA_nanopore, ecc_finder, and FLED, respectively **(Supplementary Table 2)**. 91% of eccDNAs detected by eccDNA_RCA_nanopore were overlapped with FLED-detected full-length eccDNAs, with 63% for ecc_finder, showing the reliability of FLED (**Figure 2C**). FLED displayed a significant advantage in terms of the number of detected full eccDNAs and 481 eccDNAs were found by all tools. A comparison of the length distribution of full-length eccDNAs detected by three tools also showed that FLED is more sensitive to eccDNAs with a larger size than ecc_finder and ec-cDNA_RCA_nanopore benefit from full-length reads covered the break-points only once (**Figure 2D**). Similarly, on the high-coverage simulated dataset, FLED also shows higher sensitivity and accuracy compared to the other two tools (**Supplementary Figure 2**). Collectively, since FLED made full use of reads with more than one but less than two tandem eccDNA sequence repeats, the sensitivity of FLED, especially for eccDNAs with large sizes, was greatly improved.

In addition, eccDNAs composed of multiple, non-contiguous genomic segments were verified. Two multiple-fragment eccDNAs were selected and three pairs of primers were designed for each eccDNA (**Figure 2E**). The junctions between fragments and the breakpoints of eccDNAs were validated by PCR, and the internal structures were validated by Sanger Sequencing (**Figure 2E**), demonstrating the unique structure of eccDNA different from the linear genome, while only the breakpoints were determined by Circle-Map. Taken together, these results of experimental validation demonstrated the high reliability and efficiency of FLED for eccDNA detection based on Nanopore data.

### 3.3 EccDNA detection in cell lines using nanopore sequencing datasets

Seven human cell lines from normal and tumor tissue were processed by an adapted Circle-seq protocol and sequenced using the Nanopore MinION platform, generating an average of 1M reads per sample (detailed in **Supplementary Table 3**). FLED was applied to identify eccDNAs in each sample after pre-processing including base-calling, reads quality control, and alignment (Detailed in **Supplementary Methods**). The amount of eccDNA detected by FLED in seven cell lines spanned a wide range, varying from 521 to 19326, showing the heterogeneity of cell types (detailed in **Table 1**).

**Table 1.** Nanopore sequencing and eccDNAs detected by FLED in different cell lines.

|  | BGC823 | SGC7901 | GES1 | HepG2 | HL7702 | MB453 | MCF12A |
| --- | --- | --- | --- | --- | --- | --- | --- |
| Phenotype | GC | GC | GE | HCC | Liver | BC | BE |
| Read number | 1,224,574 | 730,346 | 666,873 | 520,421 | 2,096,323 | 1,250,482 | 864,510 |
| Mapping rate | 98.36% | 98.24% | 97.93% | 98.10% | 94.76% | 92.24% | 98.40% |
| Full-length reads | 25,867<br>(2.12%) | 25,240<br>(3.47%) | 17,944<br>(2.71%) | 47,630<br>(9.30%) | 40,098<br>(2.00%) | 93,839<br>(7.96%) | 2,240<br>(0.26%) |
| Partial reads with BP | 9.765% | 9.407% | 8.615% | 11.389% | 4.022% | 7.991% | 0.725% |
| eccDNA number | 4589 | 4428 | 5780 | 5030 | 2817 | 19326 | 521 |
| Full eccDNA number | 3625 | 3651 | 4170 | 4490 | 2356 | 17388 | 323 |
| Full eccDNA with $\geq 2$ BP | 2799 | 2955 | 3077 | 3944 | 1951 | 14969 | 215 |
| Full eccDNA with 1 BP | 826 | 696 | 1093 | 546 | 405 | 2419 | 108 |
| Break eccDNA | 964 | 777 | 1610 | 540 | 461 | 1938 | 198 |
| Full eccDNA rate | 78.993% | 82.453% | 72.145% | 89.264% | 83.635% | 89.972% | 61.996% |

Note: BP: breakpoint; GC: gastric cancer; GE: gastric epithelial cells; HCC: hepatocellular carcinoma; BC: breast cancer; BE: breast epithelial cells.

Reads containing at least one complete copy of eccDNA were considered full-length reads. The proportion of full-length reads in libraries ranges from 2% to 9%. Based on these reads, the full-length sequences of an average of 75% detected eccDNAs in seven cell lines could be obtained without assembly, providing direct evidence of internal structure in eccDNAs (detailed in **Table 1, Supplementary Table 4**). These full-length sequences are necessary for sequence-based analysis subsequently such as structural variation and gene transcription potential analysis. In addition, there were about 10% of RCA long reads that only contained partial sequences of eccDNAs but crossed the joined junctions, hence the joined breakpoint coordinates of genomic regions can be determined from their split alignments accurately. However, the internal structure of eccDNA could not be determined from the breakpoint information alone. In this study, we labeled eccDNAs with full-length sequences as “full eccDNA”, while others with only breakpoint position as “break eccDNA”. The numbers of full eccDNAs in seven cell lines detected by FLED were 3625, 3651, 4170, 4490, 2356, 17388, and 323, respectively. Furthermore, the size of eccDNAs was defined as the sequence length for full eccDNAs and the distance between breakpoints for break eccDNAs, respectively. The size distributions of eccDNAs were similar among all seven cell lines, ranging from 0.06kb to 7.7Mb with a median of 2.64kb. Furthermore, a comparison was performed between the size distributions of full and break eccDNAs and showed a significant difference: The full eccDNAs have a smaller size with a median of 2.1kb while the length of break eccDNAs was heavily overestimated due to the lack of internal sequences (**Figure 3A**), suggesting that the RCA method might perform better on small eccDNAs of which full-length sequences are easier to obtain. Previous studies reported that 99% of eccDNAs are shorter than 25kb [1]. The longest full-length eccDNA detected by ecc_finder and eccDNA_RCA_nanopore was 16 kb, whereas the longest full-length eccDNA detected by FLED reached up to 30kb, and FLED identified 10 eccDNAs longer than 16 kb, indicating the improved performance of FLED for full-length eccDNA detection. However, for the possible larger eccDNA, direct detection of its full-length sequence still faces challenges due to the inefficiency of RCA. Although most detected eccDNAs are derived from a single genomic locus, FLED also found eccDNAs consisting of multiple discontinuous fragments in seven cell lines. 133 of 159 detected multiple-fragment eccDNAs included 2 or more fragments from different chromosomes, while 26 multiple-fragment eccDNAs were intra-chromosomal. Considering that these multiple-fragments eccDNAs may have independent biogenesis, they were not included in subsequent analysis.

**Figure 3.**
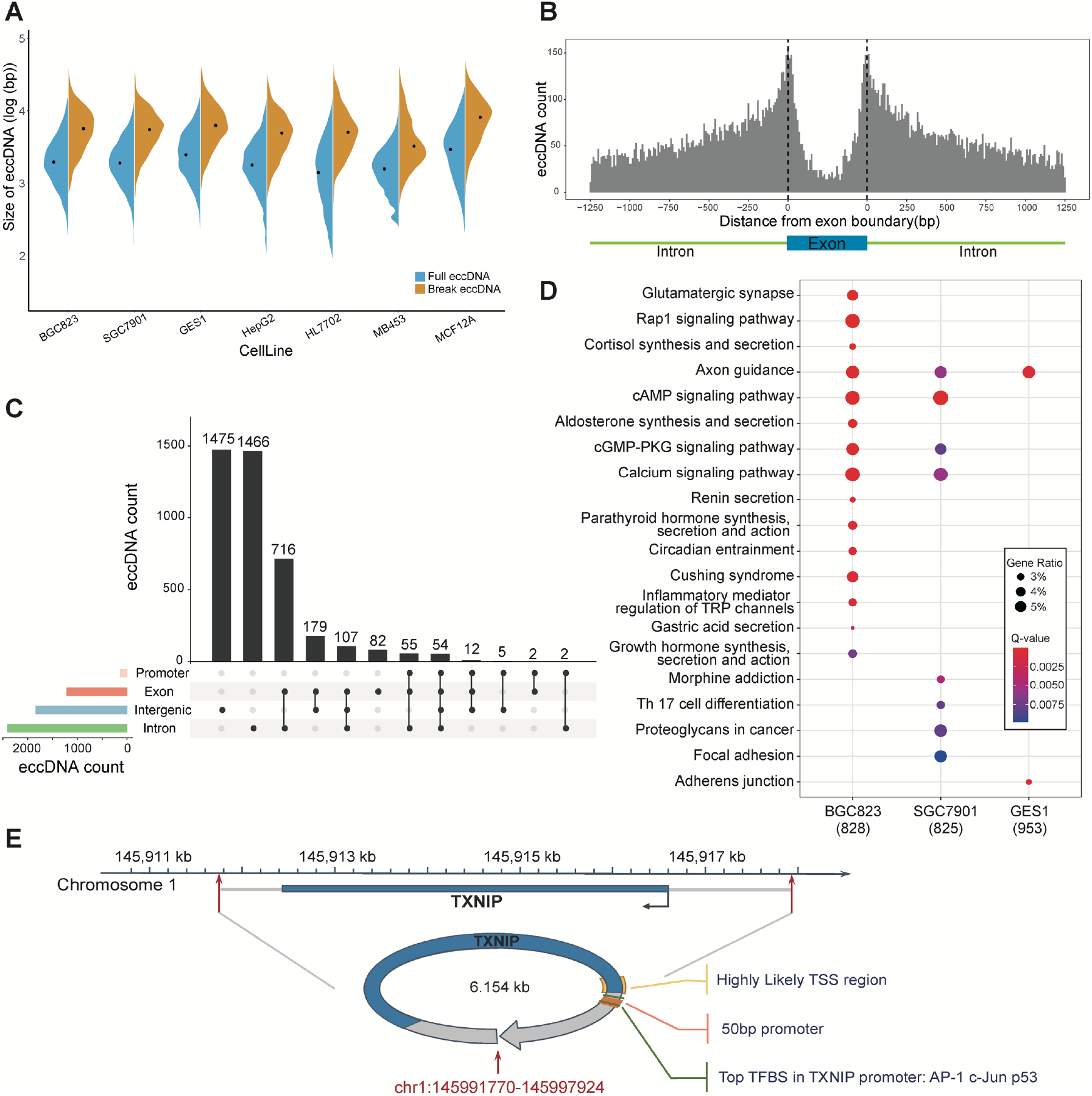
characteristics and genomic annotation of eccDNAs. **A**, The size distribution of Full eccDNAs and Break eccDNAs in seven cell lines. **B**, The distribution of Full eccDNA breakpoints around the exon-intron boundary. **C**, Genomic annotation overlap of detected Full eccDNA in GES1 cell line. **D**, KEGG pathway enriched by annotated genes in gastric-related cell lines (BGC823, SGC7901, and GES1). **E**, Display of chromosome 1 at TXNIP gene and detected eccDNA chr1:145991770-145997924, and predicted promoter (orange box), corresponding TFBS (green box), and highly likely TSS region (yellow box).

### 3.4 Characteristics and functional analysis of eccDNAs

We further analyzed the distribution and potential function of eccDNA at chromosome and gene levels. EccDNAs were founded on all chromosomes (**Supplementary Figure 3A)**, and interestingly, higher eccDNA densities appeared on both chromosomes 5 and 20 in multiple cell lines (**Supplementary Figure 3B)**. Due to the reliability of Full eccDNAs detected by FLED, subsequent analysis at the gene level was limited only to Full eccDNAs. While the eccDNA breakpoint was more enriched around exon-intron boundaries, especially occurred frequently on flanking regions of exons **(Figure 3B)**, more than half of the Full eccDNAs can be annotated to genes, and some could even contain complete genes and enhancer-like signatures, which also indicated that eccDNA might affect gene regulation through disturbing these complete genes and enhancers (**Table 2 and Figure 3C**). We applied the GO and KEGG pathway enrichment analysis for annotated genes (N=828, 825, and 953) of Full eccDNAs in three gastric-related cell lines (BGC823, SGC7901, and GES1). Although the numbers of genes enriched in BGC823 and SGC7901 (both were gastric cancer cell lines) were smaller than that in human gastric epithelial cell line GES1, more significant pathways were enriched such as Rap1 signaling pathway, Calcium signaling pathway, and cAMP signaling pathway (**Figure 3D, Supplementary Figure 4**), which were reported to be involved in the process of tumor cell migration, invasion, and metastasis in many kinds of cancers [33-35]. These pathways, which were enriched by the eccDNAs annotation genes, exerted some correlations with tumor activities, suggesting that eccDNAs might be involved in the development and progression of tumors.

**Table 2.**
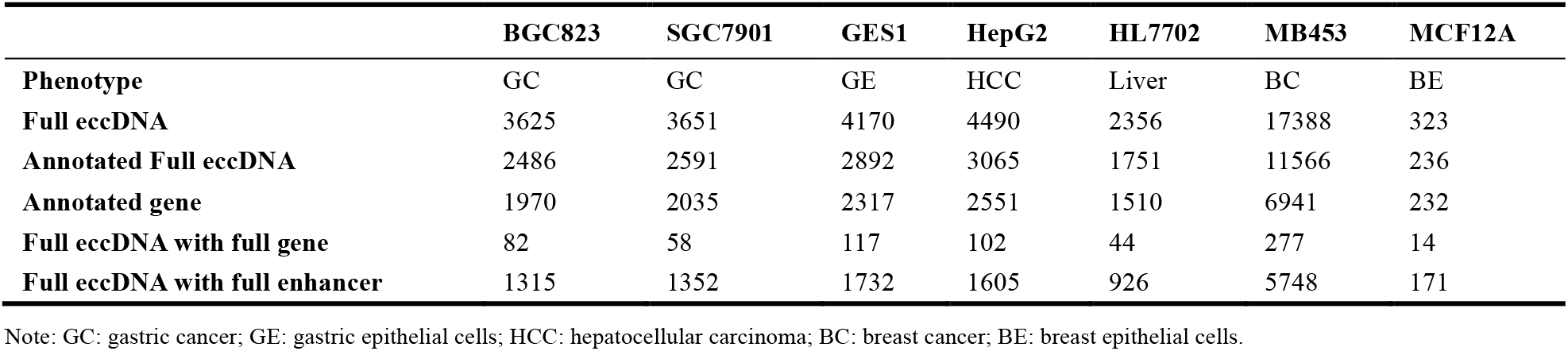
Annotation of Full eccDNA in cell lines. The number of detected Full eccDNA, annotated Full eccDNAs which overlap with gene region and Full eccDNAs contain full gene in seven cell lines respectively.

Previous studies have shown that intact genes carried by eccDNA can be expressed uncontrollably, which in turn affects transcription levels [7, 12, 13]. To investigate the impact on transcription, we performed differential expression analysis between gastric cancer cells and gastric epithelial cell GES1 based on RNA-seq, revealing that genes encoded on eccDNA, are significantly overexpressed. We identified an eccDNA named chr1:145991770-145997924 in gastric epithelial cell line GES1, which contains the complete TXNIP gene (**Figure 3E**). TXNIP was known as a tumor suppressor, which inhibits tumor cell proliferation and cell cycle progression [36-38]. The functional elements of TXNIP were predicted on eccDNA chr1:145991770-145997924, including the corresponding promoter, TFBS, and TSS. Furthermore, the upregulation of TXNIP in the gastric epithelial cell line (P-value = 0.003) was consistent with the expression pattern of tumor suppressors, indicating the transcriptional potential of eccDNAs. However, further experimental verification was needed to prove that the overexpressed TXNIP was directly transcribed from eccDNAs or genome linear DNA. Similarly, CCAT2 was encoded in eccDNA chr8:127400294-127404673 completely (**Supplementary Figure 5**), which was known as a gastric cancer-related lncRNA, involved in the differentiation and invasion of gastric cancer and has a certain prognostic ability [39, 40]. The promoters, TFBS, and TSS of gene CCAT2 were predicted on the downstream of gene, which could still play a role due to the circular nature of eccDNAs.

## 4 Discussion

In this work, we reported FLED, an experimental and bioinformatics approach for the detection of full-length eccDNA based on nanopore sequencing technology. Benefiting from the unique closed circular structure of eccDNA, eccDNA was amplified through RCA and sequenced, resulting in the multiple tandem copies of template eccDNA sequences that can be captured in ultra-long nanopore reads. FLED was assessed using simulated datasets, corresponding Illumina data, and experimental validation, demonstrating the high efficiency and reliability of our method in fulllength eccDNA detection from nanopore data.

FLED achieved a high F1 score (0.97) of eccDNA detection in the 30X simulated dataset, and 90% of 11 randomly selected eccDNA was verified by PCR and the Sanger sequencing in real data. These results proved that FLED is reliable and accurate in eccDNA detection. Moreover, while short reads-based approaches like circle-Map can identify eccDNA breakpoints according to the discordantly mapped paired-end reads and soft clipped reads, FLED detected much more full-length eccDNAs than Circle-Map, and the average size of full-length eccDNAs is 2kb, showing the advantage of FLED over the NGS-based methods. Other nanopore-based eccDNA detectors still have limitations in obtaining accurate full-length sequences of eccDNAs: Nanocircle identified the joined breakpoints only, which did not take full advantage of nanopore ultra-long reads. CReSIL introduced uncertainty in de novo assembly, and cyrcular-calling-work-flow, on the other hand, determined eccDNA according to the posterior probability. The eccDNA_RCA_nanopore and ecc_finder identified ec-cDNA from nanopore RCA reads directly but usually kept reads with at least two tandem repeats of original eccDNA sequences, which not only caused data waste but also unavailability of large-size eccDNA detection. But FLED constructed directed graphs by distinguishing the start and end location of subreads to keep as many reads as possible, which had a great advantage in the sensitivity of eccDNAs detection especially for large-size eccDNAs. The annotation of eccDNAs on the gene level also showed that eccDNAs may play an important role in the oncogenesis, metastasis, and recurrence of tumors. Some cancer-associated pathways were enriched in both gastric cancer cell lines (BGC823 and SGC7901), and it is reported that cAMP, calcium, and Rap1 signaling pathway are related to the progression of multiple kinds of cancers including gastric cancer. Moreover, the eccDNAs that contain complete CCAT2 and TXNIP genes also suggested the regulatory role of eccDNAs in cancer cell growth, migration, and invasion.

However, there is still room for improvement in both experimental and computational methods. For eccDNAs above 10kb, it is difficult to generate a mass of RCA products that comprise multiple copies of template eccDNAs, and therefore the correction of template sequences cannot perform well, which affects the accuracy of full-length sequences of eccDNAs. The proportion of full-length reads will be improved by optimizing experimental conditions or size selection. Although we avoided the identification of eccDNA templates on raw reads and used non-full-length reads for base correction to improve data utilization, full-length eccDNAs were determined based on the full-length reads, and for extremely large eccDNA, particularly, it is almost impossible for Phi29 DNA polymerase to process the eccDNAs beyond a full circle, resulting in the full-length sequences of these eccDNAs is unavailable without assembly. Moreover, limited by the current sequencing depth, FLED is far from saturation, and the whole profile of eccDNAs can be provided with deeper sequencing. In conclusion, FLED is a reliable full-length eccDNA detection tool based on nanopore sequencing data and the analysis of the detailed structure of eccDNA may contribute to the in-depth understanding of the occurrence and development mechanism of cancer.

### Key Points

- Extrachromosomal circular DNA (eccDNA) is widespread in eukaryotes and plays a role in biological processes. In the study, we developed a novel FLED method to identify eccDNA and its accurate full-length sequence for long-read sequencing data.
- In FLED, we constructed a directed graph independently for each read to ensure the accuracy of identified eccDNA, then incorporated high-quality non-full-length reads to correct sequencing errors.
- FLED could effectively identify eccDNA and its internal structure, which provided new insight into an in-depth understanding of the occurrence and development mechanism of cancer.

## Availability and requirements

The data underlying this article has been deposited at National Genomics Data Center with the accession number HRA002605 (temporary URL: https://ngdc.cncb.ac.cn/gsa-human/s/YI9346cq).

FLED is implemented using *Python3* and is freely accessible at https://github.com/FuyuLi/FLED. The scripts to generate the simulation data, along with the simulated Circle-seq datasets and the reference genomic sequences are also available on GitHub.

## Supplementary Data

Supplementary data are available online at *Briefings in Bioinformatics*.

## Funding

Y.B is supported by the National Natural Science Foundation of China (grant number: 61871121). X.D. is supported by American Parkinson’s Disease Association, ASAP, and NIH (grant U01NS120637). This research was funded in whole or in part by Aligning Science Across Parkinson’s ASAP-000301 through the Michael J. Fox Foundation for Parkinson’s Research (MJFF). For the purpose of open access, the author has applied a CC BY public copyright license to all Author Accepted Manuscripts arising from this submission. The funders had no role in the study design, data collection, analysis, the decision to publish, or the preparation of the manuscript.

## Acknowledgments

Not applicable.

## Author contributions

This study was conceptualized and designed by Yunfei Bai, with co-supervision and consultancy from Xianjun Dong. The experiment and Nanopore sequencing were performed by Wenxiang Lu. The tool development and data analysis were completed by Fuyu Li. Fuyu Li, Wenxiang Lu, Wenlong Ming, and Xianjun Dong wrote the manuscript with input from all other authors. All authors read and approved the submission and publication.

## Competing interests

The authors declare that they have no competing interests.

## Notes

### Competing Interest Statement

The authors have declared no competing interest.

