## Supplementary Figures for "Detecting Full-Length EccDNA with FLED and long-reads sequencing"

Supplementary Methods, Figures and Tables

### Supplementary Methods

#### Cells culture and DNA extraction.

GES1, SGC7901, BGC823, HepG2, HL7702, MCF12A, and MD-MB453 cells were grown in Dulbecco’s Modified Eagle’s Medium (DMEM, GIBCO), which were all supplemented with 10% fetal bovine serum (FBS, GIBCO) with 100 μg/mL streptomycin and 100 units/mL penicillin at 37°C in 5% CO2. DNA extraction was performed using a Blood & Cell Culture DNA Kits (Qiagen) and quantified using Qubit dsDNA High Sensitivity Kit (Invitrogen).

#### Circle-seq for eccDNAs enrichment

The genomic DNA was subjected to Plasmid-Safe ATP-dependent DNase (Epicentre) digestion at 37 °C for 24 hours with a final concentration of 0.4 U/µl to remove linear dsDNA. To ensure complete hydrolysis of linear dsDNA, additional ATP and DNase (0.2U/µl) were added every 24 h and perform 5 days.

For eccDNA amplification, each eccDNA-enriched sample was performed random rolling circle amplification according to previous reports [1]. A 20 µl reaction was set up using 1 mM dNTPs, 10 U Phi29 DNA polymerase (NEB), 50 µM Exo-resistant random primer (Thermo Fisher Scientific), 0.02 U inorganic pyrophosphatase (Thermo Fisher Scientific) and 1× Phi29 DNA polymerase buffer. The reaction was run at 30 °C for 18 h and stopped by heating to 65 °C for 2 min. Product DNA was purified by AMPure XP beads.

#### Nanopore library generation and sequencing

For nanopore sequencing libraries preparation, the de-branching reaction of Phi29-amplified DNA was required. In accordance with instructions provided with T7 Endonuclease I, 2.5 µl of T7 Endonuclease I was added to the reaction, and incubated at 37 °C for 1 h, followed by purification through AMPure XP beads. The de-branched DNA was used to prepare libraries for nanopore sequencing using the Ligation Sequencing Kit LSK109 (Oxford Nanopore), and then performed on a MinION instrument to sequencing according to the manufacturer’s protocol (Oxford Nanopore, flow cell R9.4.1).

#### Base-calling, quality control, and alignment of nanopore sequencing datasets

The raw nanopore current signal trace was recorded in FAST5 format, and GPU base-calling, demultiplexing as well as barcode trimming were performed using the Guppy base-caller and barcoder version 2.3 in the R9.4 high-accuracy model. The cleaned reads with quality below 7 were discarded by NanoFilt. High-quality reads were aligned to the human reference genome (GRCh38) to record the origin of chromosomal derived eccDNAs using minimap2 (version 2.17-r941) with standard ONT parameters, which was specifically designed for long noisy reads alignment and supported spliced alignment.

#### Simulation-based validation

NanoSim (Yang, et al., 2017) was used to generate simulated nanopore sequencing datasets. The alignment file of GES1 reads was provided to characterize the experimental MinION sequence data firstly, and its sequencing errors and length distribution were featured serving as the input in the simulation step. Based on the detected eccDNA list from GES1 cell line data, the full-length sequences of simulated eccDNA were extracted from the reference genome according to breakpoint coordinates and tandemly repeated 10 times to simulate the RCA products.

Using the read model built in the previous step, 2 datasets with different sequencing depth were simulated; 200000 reads from the canonical reference genome and 200000 reads from 4297 simulated eccDNA were generated and pooled together reaching a sequencing depth of 30x. Another lower depth (10x) dataset was also simulated from 4153 eccDNA.

The simulated datasets were filtered and mapped with the same criteria as real reads, then eccDNA was identified by FLED.

To evaluate the performance of FLED, the sensitivity, defined as the number of detected eccDNA present in the simulation set, and the precision, defined as the fraction of correctly identified eccDNA found on the simulated set were calculated. A single metric F_1_ score was assessed, which equally favors an increase in sensitivity and precision as the harmonic mean of the sensitivity and precision, according to the formula:

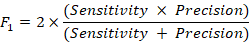

#### EccDNA annotation

Coordinates of Full eccDNAs obtained by FLED was intersected using *bedtools intersect* version 2.29.2 (Quinlan and Hall, 2010) with coordinates of genes to determine the genes overlapped with eccDNAs. The gene coordinates were extracted from the evidence-based annotation of the human genome (GRCh38), version 27 (Ensembl 90) downloaded from GENCODE. For full eccDNAs without gene overlapped, ChIPseeker (Yu, et al., 2015) was performed to retrieve the eccDNA located on the upstream (3kb) of genes.

The ENCODE Registry of candidate cis-Regulatory Elements (cCREs), including promoter-like signature and enhancer-like signature, are downloaded from UCSC download server, and only enhancers that are fully contained in the eccDNA regions can be annotated.

BDGP was used to identify the promoter region based on the sequences of eccDNA, and Promoter2.0 was applied to identify the likely TSS with only highly likely TSS was considered. For TFBS, PROMO constructs specific binding site weight matrices for the TFBS defined in TRANSFAC database to identify putative TFBS and the TFBS recorded in GeneCards were remained.

#### Illumina library generation and sequencing

Libraries for next-generation sequencing were prepared using the Hieff NGS OnePot DNA Library Prep Kit for Illumina (Yeasen) according to the manufacturer’s protocol. Libraries were sequenced on NovaSeq instruments with 2 × 150 bp paired-end reads.

#### EccDNA detection in Illumina dataset

Quality control of Paired-end Illumina sequencing data was performed by FastQC, and the clean reads were aligned to the human reference genome (GRCh38) by BWA-MEM with default settings. Circle-map was used to identify eccDNA with the same filtering criteria as FLED.

#### EccDNA detection by other ONT-based eccDNA detector

The eccDNA_RCA_nanopore script was download from <https://github.com/icebert/eccDNA_RCA_nanopore> and the alignment was completed by minimap2 with parameters of “-x map-ont -c --secondary=no”. For ecc_finder (version v1.0.0, https://github.com/njaupan/ecc_finder), the long-read-mapping mode was applied with parameters of “--min-read 1 --min-bound 0.5 --min-cov 2”. The candidate eccDNAs supported by at least 3 distinct nanopore reads were reserved for comparison.

### Supplementary Figures

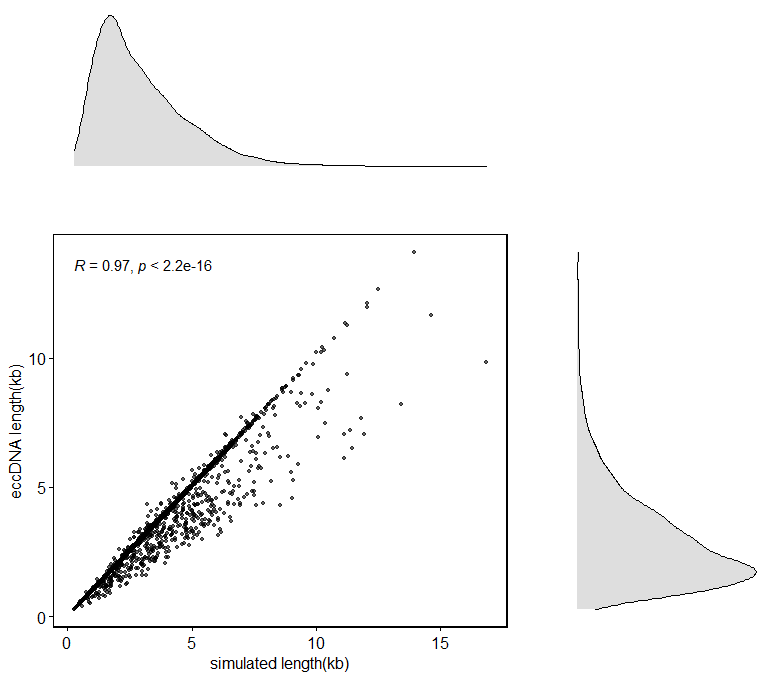

**Supplementary Figure 1:** The length distribution of detected eccDNAs on the 30x simulated dataset generated by NanoSim, showing a high correlation (Pearson’s r = 0.9, p-value of 2.2
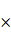
10^-26^) especially for eccDNAs with small size.

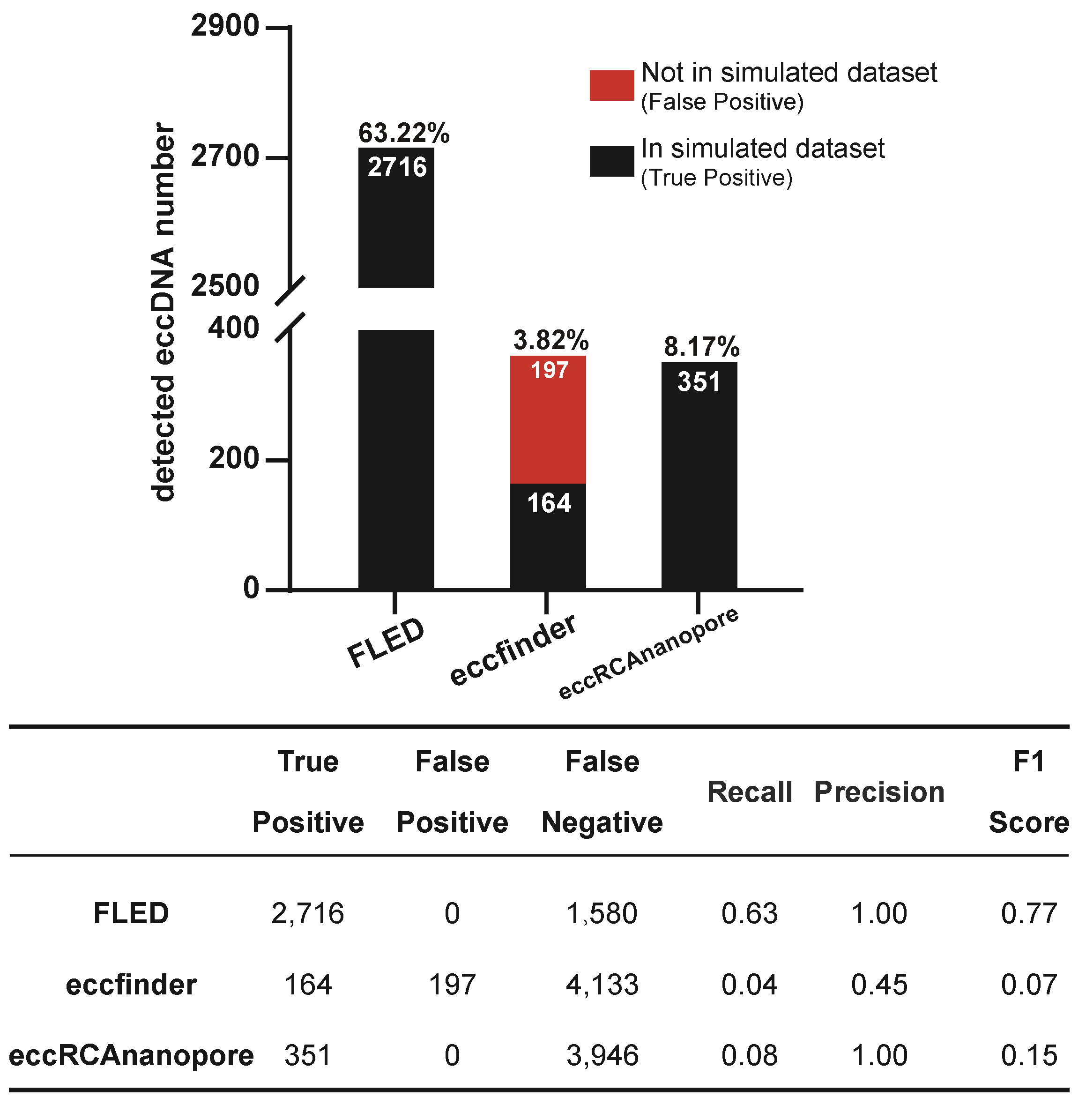

**Supplementary Figure 2:** The number of full-length eccDNAs detected by FLED (2,716 in total, Recall: 63.22%), ecc_finder (361, 3.82%) and eccDNA_RCA_nanopore (351, 8.17%) in high-coverage simulated dataset.

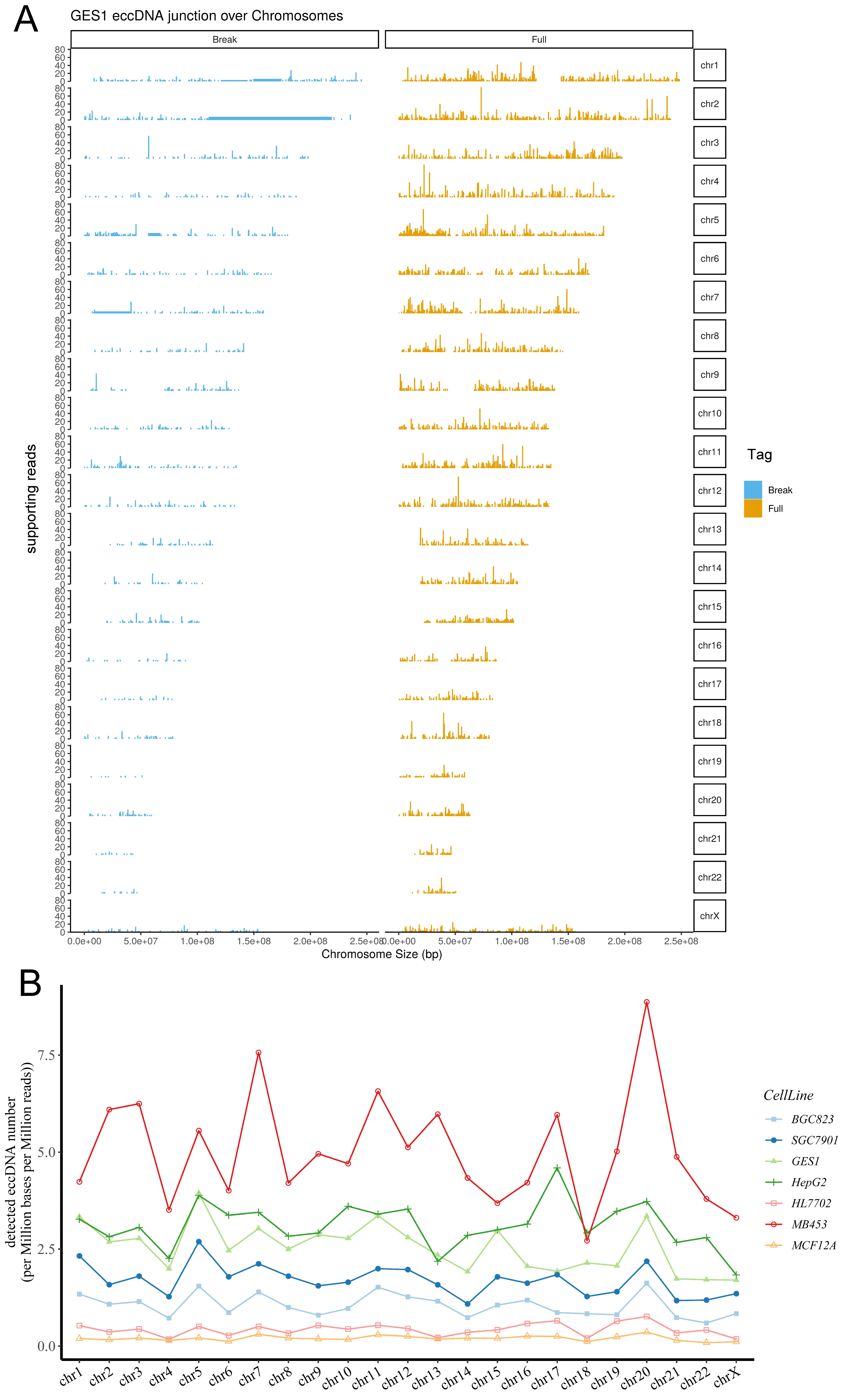

**Supplementary Figure 3:** genomic distribution of eccDNAs. **A,** Distribution and supporting reads number of detected eccDNAs (Full and Break) breakpoints across the human genome in GES1 cell line. **B,** EccDNAs frequency relative to chromosome in seven cell lines. Full: eccDNAs with full-length sequences obtained; Break: eccDNAs with only breakpoint position were obtained.

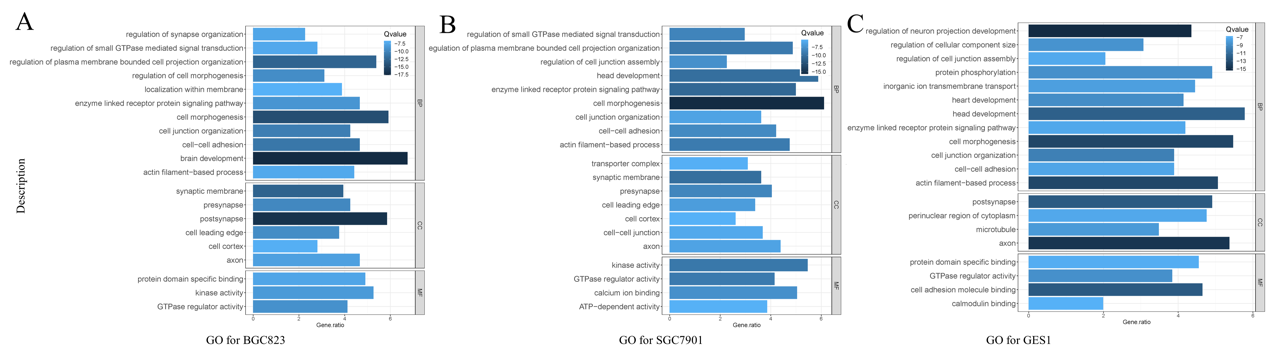

**Supplementary Figure 4:** The GO enrichment analysis of eccDNA-annotated genes in three gastric cell lines: gastric cancer cells BGC823 and SGC7901 and gastric epithelial cells GES1.

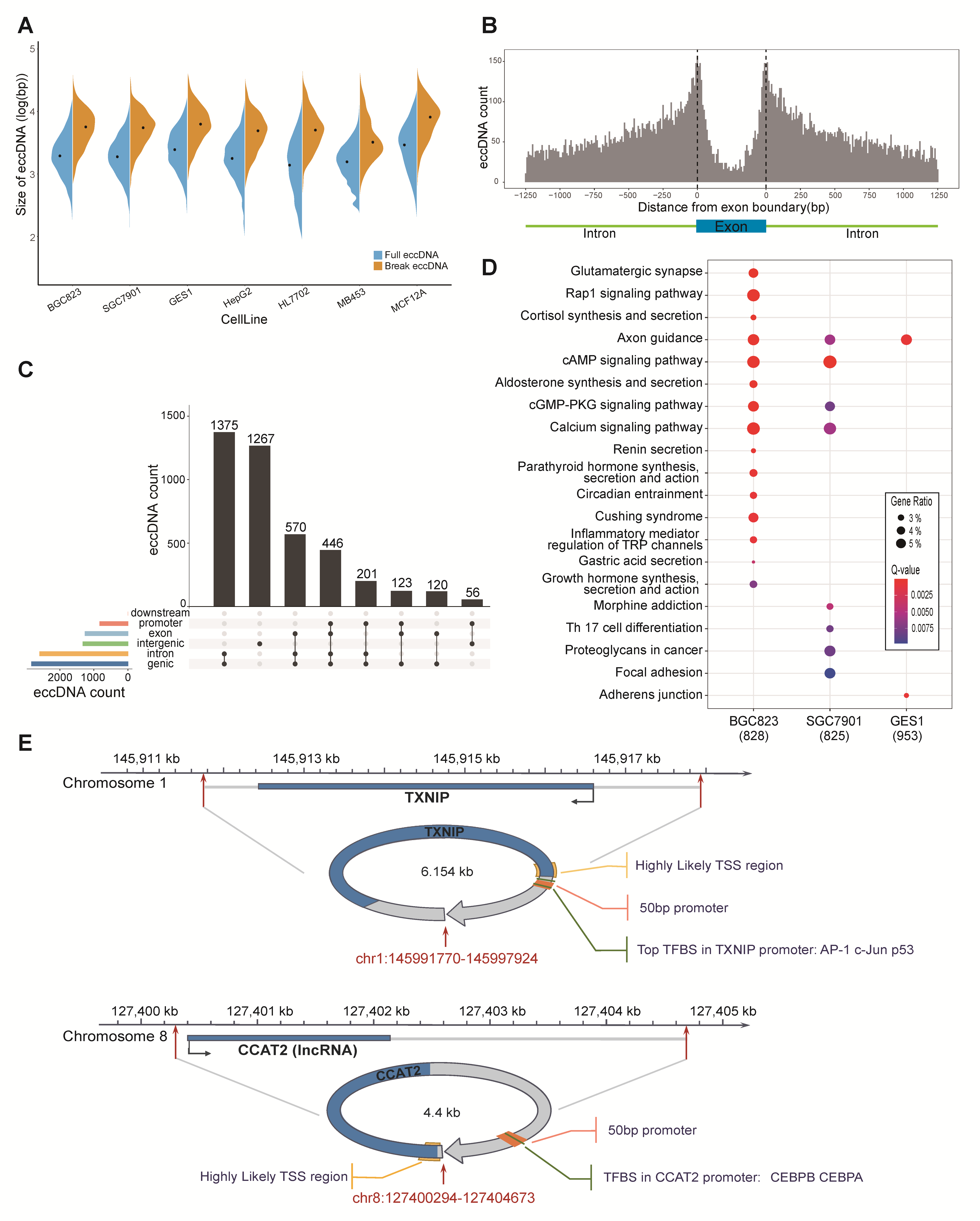

**Supplementary Figure 5:** Display of chromosome 8 at CCAT2 gene and detected eccDNA chr8:127400294-127404673, and predicted promoter (orange box), corresponding TFBS (green box), and highly likely TSS region (yellow box).

### Supplementary Tables

**Supplementary Table1. The PCR primers for eccDNA validation.**

**Supplementary Table2. List of detected eccDNAs by three nanopore-based detectors in GES1 cell line.**

**Supplementary Table3. Summary of data produced by Nanopore sequencing.**

**Supplementary Table4. List of detected eccDNAs by FLED in seven cell lines.**
